# Fertility recovery after combined adriamycin, bleomycin, vinblastine and dacarbazine chemotherapy in mice

**DOI:** 10.1101/2025.05.01.651671

**Authors:** Xinmei Lu, Bongkoch Turathum, Ri-Cheng Chian, Yubing Liu

## Abstract

**Background:** With combined adriamycin, bleomycin, vinblastine and dacarbazine (ABVD) chemotherapy and radiotherapy, most of Hodgkin lymphoma (HL) patients were cured. The effect of ABVD treatment on ovarian reserve and fertility has not been fully characterized, and whether fertility preservation is required is controversial and not empirically known.

**Methods:** Eight-week-old female mice were injected intravenously with saline solution or ABVD for 4 weeks, respectively. Estrous cycles, level of serum anti-Mullerian hormone (AMH), body weight and ovary weight were recorded. Ovarian tissue structure and the number of follicles at each stage were analysed. IVF assay and natural mating trials were performed. Finally, transcriptome sequencing of ovarian tissue was conducted.

**Results:** Most mice completely lost their estrous cycle soon after ABVD injection. After 4 cycles of treatments, the body weight gain and gross ovarian weight were significantly lower than those of the control. However, estrous cycles were returned to normal at 4 weeks after ABVD discontinuation. Serum AMH and the numbers of primordial follicles were significantly increased compared to controls. ABVD withdrawal did not affect the rate of cleavage or blastocyst formation in IVF assays. Furthermore, the mice were fertile and there was no significant difference in offspring weight, but the number of offspring in the ABVD discontinuation group was smaller compared with the controls. Transcriptome analyses revealed that the JAK/STAT pathway, which is involved in the inhibition of primordial follicles activation and apoptosis, was upregulated in ABVD-treated ovaries, whereas few gene expressions were altered 4 weeks after ABVD withdrawal.

**Conclusions:** The ovarian reserve and reproductive function can be recovered after ABVD treatment, and combined ABVD treatment for a short time has a beneficial effect on the ovarian reserve, which can be used to provide a basis for guiding fertility preservation decision-making for HL patients.

**Statements and Declarations:** The authors declare no competing interests.

## Background

Hodgkin lymphoma (HL) is a B-cell haematological malignancy that usually emerges before adolescence and adulthood[1]. Rapid improvements in the survival rate of HL treatment in recent decades have made it one of the most curable human cancers[2]. At present, the adriamycin, bleomycin, vinblastine, and dacarbazine (ABVD) regimen is the most common first-line treatment of HL in worldwide and is considered to be the “gold standard” chemotherapy for HL[3, 4]. With combined chemotherapy and radiation therapy, more than 80% of patients were cured[5]. However, there is now more concern regarding the long-term side effects of anticancer treatments, and female fertility status is a prominent issue[6–8].

Female fertility is based on the pool of primordial follicles in the ovaries (*i.e.,* the ovarian reserve). The size of the pool of primordial follicles progressively decreases with age. For women of reproductive age, chemotherapy-induced gonadotoxicity may accelerate this loss, leading to diminished ovarian reserve (DOR) and even premature ovarian failure (POF) with loss of fertility in female survivors of cancer, which is usually described as chemotherapy-induced diminished ovarian reserve (chDOR) or chemotherapy-induced premature ovarian failure (chPOF)[9–11].

The type of chemotherapeutic agent affects the level of ovarian damage or fertility loss. The majority of publications have reported that ovarian damage is mainly caused by alkylating agents[12]. There are reports verifying that advanced-stage HL patients treated with one or more alkylating agents have a higher prevalence of ovarian dysfunction and reduced fertility potential[13]. In contrast, there are reports suggesting that in early-stage HL patients receiving ABVD treatment, that the ovaries have a good reproductive prognosis in terms of chDOR and chPOF[14]. There are also studies showing that ABVD treatment has a lower risk of significant fertility impairment or POF.

ABVD causes no irreversible impairment of female reproductive functionk[15, 16]. Female HL patients who had survived without recurrence for ≥3 years and who had attempted pregnancy after ABVD chemotherapy did not experience significant subfertility[15]. The risk of amenorrhea before the age of 40 was not increased in ABVD chemotherapy patients[14]. It has been reported that the levels of AMH returned to premedication levels 12 months after ABVD treatment ended[17, 18]. A recent study even demonstrated that ABVD treatment does not diminish the ovarian reserve and may paradoxically increase the number of primordial follicles[19]. In contrast, it has also been reported that treatment with ABVD can lead to irreversible damage to female fertility, requiring fertility preservation, but the specific mechanism is unclear[8].

Although ABVD is considered a treatment with a low gonadotoxic risk, the published studies have been limited by a small study population and heterogeneity and some studies have reported that HL itself affects ovarian function. Thus, there is still controversy as to whether fertility preservation should occur before ABVD treatment. Presently, the effect of ABVD treatment on the ovarian reserve and fertility in women of childbearing age has yet to be fully characterized. Additionally, whether fertility preservation is required is controversial and not empirically known. Here, we systematically investigated the effects of ABVD on reproductive function using a mouse model. Our findings have valuable information with fertility preservation guidance for HL patients.

## Methods

### Animals and experiment design

All mouse experimental procedures were approved by the Ethics Committee of Shanghai Tenth People’s Hospital and conducted in accordance with Tongji University animal research requirements. ICR female mice (6 weeks old) and male mice (8 weeks old) were purchased from Beijing Vital River Experimental Animals Centre (Beijing, China). They were raised under controlled illumination (12 h light-dark cycle) and temperature (20–25°C) conditions and fed a regular diet throughout the study period. Except for mice in mating trials, the other mice were housed with up to five mice per cage.

The experimental design is shown in Fig. 1A. The doses and treatment cycles for ABVD were chosen based on the standard dose given to humans[20, 21], calculated according to the formula: animal equivalent dose (mg/kg) = human dose (mg/kg) x km ratio, where km equals 12.3 for the mouse[22]. Briefly, 8-week-old female mice were intravenously administered saline solution or ABVD [adriamycin (8.3 mg/kg), bleomycin (3.3 mg/kg), vinblastine (2.0 mg/kg) and dacarbazine (124.7 mg/kg)] once weekly for 4 weeks. ABVD was suspended in 0.9% NaCl. The group intravenously injected with 0.9% NaCl was designated as a control group. The estrous cycle was monitored continuously after first ABVD or 0.9% NaCl injection. Animals were ultimately sacrificed at the following time points to analyse ovarian functions: finishing ABVD treatment (12 weeks old) and returning to normal estrous cycles (4 weeks after removing ABVD, 16 weeks old). Natural mating trials and in vitro fertilization assay were performed to assess fertility when estrous cycles resumed (16 weeks old). The mice were humanely treated during ovary and epididymis collection, and pain relief was considered.

**Fig. 1.**
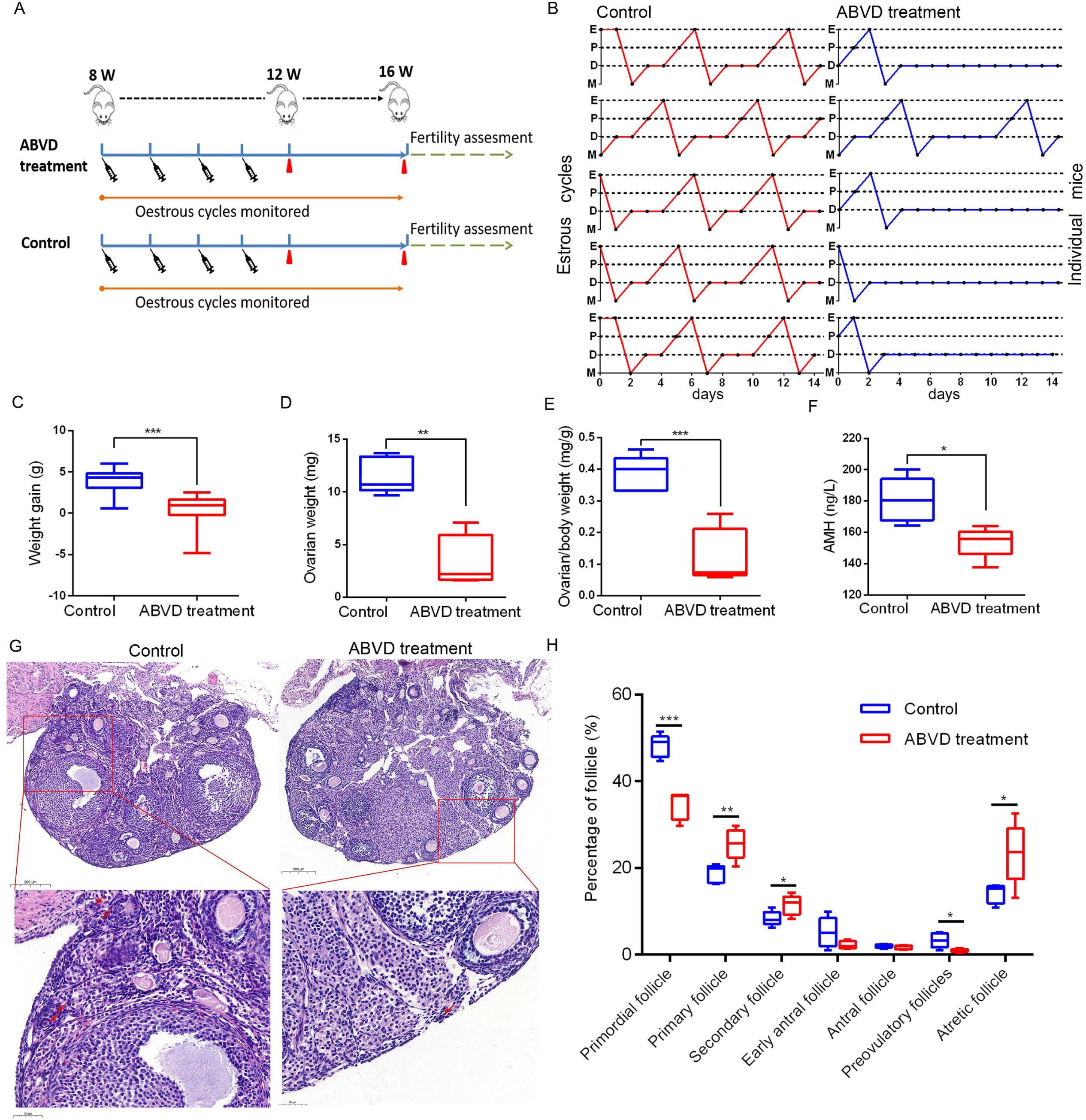
Ovarian function was disrupted after ABVD treatment. A. Schematic diagram of the experimental procedure. Female mice (8 weeks old) were injected intravenously with saline solution or ABVD once weekly for 4 weeks. Specimens were collected at the times of ABVD treatment completion (12 weeks old) and estrous cycles resumed (16 weeks old) respectively for ovarian function assessment. Fertility was assessed when estrous cycles resumed (16 weeks old). Red-filled triangles indicate sample collection time points. Each syringe represents a saline solution or ABVD injection. B. Estrous cycles were monitored by vaginal smear assays daily for 14 days after ABVD treatment (n = 5 per group). M, metestrus; D, diestrus; P, proestrus; O, oestrus. C-F. Comparison of weight gain, ovarian weight, ovary organ index and serum AMH concentrations between the ABVD treatment and control groups. G. Representative histology of ovary sections (H&E staining). Red arrow, primordial follicles. H. Distribution of follicles at different stages in each ovary (n = 5). The data are presented as the means ± SDs. \**P* < 0.05, \*\**P* < 0.01 and \*\*\**P* < 0.01, compared with the control group.

### Estrous cycle determination

After ABVD treatment, the estrous cycle stage was determined by consecutive vaginal smears[23]. Briefly, vaginal secretions were collected daily (9:00 am) with saline (0.9% NaCl), spotted onto a dry microscope glass slide, subjected to haematoxylin-eosin staining for 3–4 min after air-drying at room temperature, washed and dried. The stage of the estrous cycle (proestrus, estrus, metestrus, and diestrus) (4–5 days total) was determined by cytology under a light microscope, as previously described[24].

### Hormone measurements

For the serum hormone analysis, orbital blood of mice was collected into 1.5-mL tubes and placed at room temperature for 1 h to coagulate spontaneously. Serum was then separated by centrifugation at 3000 rpm for 10 min at 4°C. The level of serum anti-Mullerian hormone (AMH) was measured with an enzyme-linked immune sorbent assay (ELISA) kit (SenBeiJia Biological Technology, Nanjing, China) according to the manufacturer’s instructions[25].

### Histological analysis of ovaries

The mice were anaesthetized with urethane (0.6 mL/100 g), the adipose tissue was removed by peeling, and the ovaries were collected from each group (ABVD treatment and control female mice). The left ovary of each mouse was fixed in 4% (w/v) paraformaldehyde (pH 7.5) at 4°C overnight, dehydrated and embedded in paraffin. The right ovary of each mouse was snap frozen in liquid nitrogen and stored at -80°C. The ABVD withdrawal mice and their respective controls were euthanized at diestrus. Then, the paraffin-embedded ovaries were sectioned into 5 μm-thick slices for haematoxylin and eosin (H&E) staining.

The morphology of ovarian follicles was observed under a light microscope (Nikon; Tokyo, Japan) at 400x magnification. We examined the distribution of ovarian follicles at different developmental stages. Briefly, follicles with an oocyte surrounded by a single layer of squamous or cuboidal granulosa cells were classified as primordial follicles and primary follicles; follicles with two or more layers of cuboidal granulosa cells with no visible antrum were classified as secondary follicles; Early antral follicles with four or more layers of granulosa cells and an antrum diameter < 20 μm, whilst antral follicles contain a single clearly defined antral space; Follicles with the largest antral space and a defined cumulus granulosa cell layer were classified as preovulatory follicles; Follicles containing a degenerating oocyte, pyknotic nuclei, shrunken granulosa cells, or apoptotic bodies were considered atretic[26, 27]. Different types of follicles of the entire ovary were quantified by stereological follicle counts methods according to published methods[28, 29]. Briefly, the entire paraffin-embedded ovaries were sectioned serially into 5 μm-thick slices. Every fifth section was selected for H&E staining and used for follicle counting. Only follicles containing oocytes with obvious visible nuclei were counted to avoid counting any follicle twice, which were classified according to their development stages into primordial, primary, secondary, antral, and atretic follicles, as previously described. Accordingly, the number of different types of follicles in the selected sections was by multiplied 5 to calculate the total number of follicles per ovary for the fact that every fifth section was used in the analysis.

### Fertility assessment

To evaluate the fertility of mice, natural mating trials were undertaken when the estrous cycles recovered after the end of ABVD treatment. Female mice from each group (ABVD treatment or control, 16 weeks old) were mated with adult males (10 to 12 weeks old) with proven fertility. Two female mice were housed in a cage with one adult male mouse, and successful mating was identified by the presence of a vaginal plug the following morning. Then, the female mice were raised in single cages until the birth of their pups. The number of delivered pups and pup weight at postnatal day 2 were recorded.

### Oocyte collection and in vitro fertilization

In vitro fertilization (IVF) and embryo culture were performed as previously described[30]. Briefly, 16-week-old female mice were given 5 IU pregnant mare serum gonadotropin (PMSG, Ningbo Hormone Product Co., China) intraperitoneally, and after 48 h, an additional injection of 5 IU of human chorionic gonadotrophin (hCG, Ningbo Hormone Product Co., China) was administered to induce oocyte maturation.

Oviductal ampullae were removed at 16 h after hCG injection and broken with an insulin syringe (1 mL) with a 31G needle (BD Biosciences, San Jose, CA) to release the cumulus oocyte complexes (COCs). After washing three times in HEPES-buffered medium, the COCs were collected in KSOM medium (EMD Millipore Corp, Billerica, MA, USA) in a 5% CO_2_, 5% O_2_ atmosphere at 37 ℃ for IVF.

Sperm were harvested from 10-to 12-week-old ICR male mice. The cauda epididymis was placed in a dish of human tubal fluid (HTF) medium to release sperm, followed by capacitation for 1 h (37°C, 5% CO_2_, and 5% O_2_). Then, sperm (1 × 10^6^/mL) were added to oocyte droplets in HTF medium for 4-6 h. The presence of two pronuclei was considered to indicate successful fertilization. Zygotes were cultured in droplets of KSOM medium covered with mineral oil incubated under 5% CO2, 5% O2, and 90% N2 at 37°C. Subsequent on-time embryonic development was assessed.

### RNA extraction and transcriptome sequencing

Transcriptome sequencing was performed by Applied Protein Technology, Ltd. (Shanghai, China). Briefly, total RNA was isolated from mouse ovaries using TRIzol Reagent (Invitrogen, USA) following the manufacturer’s instructions. RNA quantity and integrity were quantified using a NanoDrop ND-2000 spectrophotometer (Agilent Inc., USA) and an Agilent 2100 Bioanalyzer with RNA 6000 Nano Kits (Agilent Technologies, CA, USA), respectively.

A transcriptome sequencing library was constructed using rRNA-depleted RNAs with the NEBNext® Ultra™ II RNA Library Prep Kit (NEB, USA) following the manufacturer’s recommendations. Libraries were quantified using a Qubit 2.0 instrument, and library size was measured using an Agilent 2100 Bioanalyzer (Agilent Technologies, USA). The libraries were then pooled and sequenced on the Illumina HiSeq2500 system (Illumina, California, USA), generating paired-end 50 base-pair reads.

### KEGG pathway enrichment analysis

Differential expression analysis of the ABVD group and the control group was performed using the DESeq2 R package (1.16.1). The false discovery rate (FDR) was obtained using the Benjamini–Hochberg method to correct for multiple hypothesis testing. Genes with an adjusted FDR <0.05 and at least a 2-fold change by DESeq2 were considered differentially expressed. Kyoto Encyclopedia of Genes and Genomes (KEGG) pathway analysis was performed on the DEGs to analyse candidate DEG functions. The significance of KEGG term enrichment was estimated with FDR < 0.05.

### Statistical analyses

Quantitative data are expressed as the mean ± standard deviation (SD). The statistical analyses were performed using SPSS 17 (Version 22, SPSS Inc., Chicago, IL, USA) and were performed using a two-tailed unpaired Student’s t test or the Mann–Whitney U test if the data were not normally distributed. The chi-square test was used for values presented as frequencies and percentages. Differences were considered significant when *P* < 0.05.

## Results

### Assessment of the effects of 4 cycles ABVD treatment on ovarian function in female mice

As shown in Fig. 1A, we first established an ABVD treatment model with weekly intravenous injection of ABVD for 4 weeks, and the control group was treated with normal saline. Mice from the control group exhibited regular estrous cycles with a duration of 4-5 days. However, most mice lost their estrous cycle completely soon after ABVD treatment (Fig. 1B). Results showed that the total weight gain of ABVD-treated mice was significant decreased after 4 cycles treatment in comparison with control group (0.42 ± 2.05 vs. 3.95 ± 1.52 g, *P* < 0.001) (Fig. 1C). The gross ovarian weight of the control group was approximately 3.4 times greater than that of the ABVD-treated group (11.54 ± 1.72 vs. 3.48 ± 2.41 mg, *P* < 0.01) (Fig. 1D). And the ovary organ index (ovarian weight/bodyweight) was reduced significantly in ABVD-treated mice (0.13 ± 0.09 vs. 0.39 ± 0.06 mg/g, *P* < 0.001) (Fig. 1E). Additionally, serum AMH levels decreased dramatically compared with those of the control after 4 cycles of ABVD treatment (153.83 ± 9.79 vs. 180.86 ± 14.07 ng/L, *P* < 0.05) (Fig. 1F), which is similar to observations reported after ABVD treatment in women[17].

To test the effect of ABVD treatment on ovarian follicular reserve, the harvested ovaries from control and ABVD-treated mice were then fixed for morphological evaluation of follicular development. The results showed that primordial follicles appeared in clusters in the cortex of the ovary in control mice but became sporadic in ABVD-treated mice (Fig. 1G). Follicles at each stage were counted in both groups. The results showed that there was a significant decrease in the numbers of primordial follicles, antral follicles, preovulatory follicles and total follicles. In particular, ABVD treatment decreased the number of primordial follicles by approximately 61.9% (363.4 ± 60.0 vs. 138.3 ± 17.9, *P* < 0.01), while the total number of follicles decreased by approximately 46.7% (753.1 ± 109.5 vs. 401.1 ± 42.6, *P* < 0.01) in ABVD-treated mice (Table 1). The proportion of primordial follicles also decreased significantly in the ABVD treatment group (48.2 ± 2.6% vs. 34.5 ± 3.2%, *P* < 0.001) (Fig. 1H).

**Table 1.** Comparison of the number of follicles at each stage after ABVD treatment.

| Stage | Control | ABVD treatment | *P* value |
| --- | --- | --- | --- |
| Primordial follicle | 363.4 ± 60.0 | 138.3 ± 17.9 | <0.01 |
| Primary follicle | 142.5 ± 33.3 | 103.6 ± 24.1 | 0.07 |
| Secondary follicle | 61.7 ± 5.1 | 46.4 ± 12.8 | <0.05 |
| Early antral follicle | 36.2 ± 21.1 | 9.4 ± 4.6 | <0.05 |
| Antral follicle | 15.3 ± 3.5 | 6.9 ± 1.5 | <0.01 |
| Preovulatory follicle | 26.4 ± 16.0 | 4.1 ± 1.6 | <0.05 |
| Atretic follicle | 107.6 ± 28.7 | 92.5 ± 25.6 | 0.41 |
| Total | 753.1 ± 109.5 | 401.1 ± 42.6 | <0.01 |

The data are presented as the mean ± standard deviation (SD).
\* $P < 0.05$ was considered to indicate statistical significance.

### The effect of ABVD on the reproductive function of mice can be reversed

To test reproductive function after ABVD treatment withdrawal, estrous cycles were monitored daily. The results showed that the estrous cycle returned to normal 4 weeks after ABVD removal (Fig. 2A). When the estrous cycles returned to normal, there were no differences in weight gain (2.70 ± 0.85 vs. 2.04 ± 1.34 g, *P*=0.38), gross ovarian weight (7.90 ± 2.70 vs. 7.40 ± 3.00 mg, *P* = 0.79) or ovary organ index (0.26 ± 0.09 vs. 0.28 ± 0.12 mg/g, *P* = 0.85) compared with those of the control group (Fig. 2B, 2C and 2D). Interestingly, clusters of primordial follicles were clearly found in the ovarian cortex of the ABVD withdrawal mice, but fewer primordial follicles were observed in the control mice (16 weeks old) (Fig. 2F). Further analyses found that serum AMH (170.7 ± 16.6 vs. 215.7 ± 5.3 ng/L, *P* < 0.01) and the numbers of primordial follicles (121.5 ± 15.1 vs. 364.1 ± 133.1, *P* < 0.01), primary follicles (130.7 ± 18.1 vs. 312.4 ± 101.5, *P* < 0.01) and total follicles (527.6 ± 22.3 vs. 913.1 ± 269.2, *P* < 0.05) were significantly increased in the ABVD withdrawal mice (Fig. 2E, S1 and Table 2). The proportions of primordial follicles and primary follicles increased significantly in ABVD withdrawal mice, while the proportions of secondary follicles, early antral follicles, and atretic follicles decreased significantly in the ABVD withdrawal group (Fig. 2G).

**Fig. 2.**
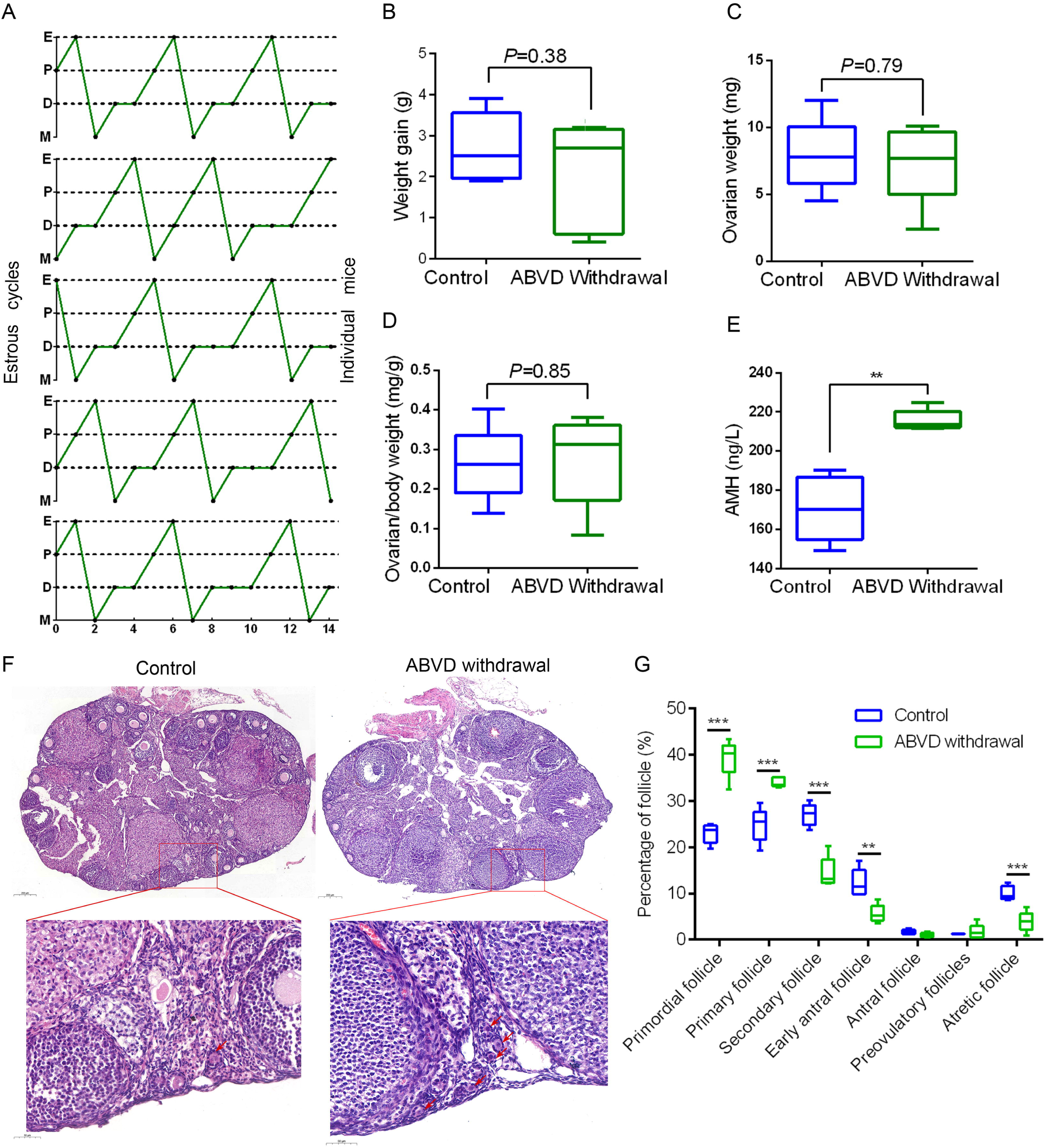
Ovarian function recovered after ABVD treatment withdrawal. A. The irregular estrous cycle recovered to normal 4 weeks after ABVD treatment withdrawal (n = 5). M, metestrus; D, diestrus; P, proestrus; O, oestrus. B-E. Comparison of weight gain, ovarian weight (from the time of saline solution or ABVD withdrawal), ovary organ index and serum AMH concentrations between the ABVD treatment withdrawal and control groups. F. Representative histology of ovary sections (H&E staining). Red arrow, primordial follicles. G. Distribution of follicles at different stages in each ovary (n = 5). The data are presented as the means ± SDs. \**P* < 0.05, \*\**P* < 0.01 and \*\*\**P* < 0.01, compared with the control group.

**Table 2.** Comparison of the number of follicles at each stage after ABVD withdrawal.

| Stage | Control | ABVD withdrawal | *P* value |
| --- | --- | --- | --- |
| Primordial follicle | 121.5 ± 15.1 | 364.1 ± 133.1 | <0.01 |
| Primary follicle | 130.7 ± 18.1 | 312.4 ± 101.5 | <0.01 |
| Secondary follicle | 142.3 ± 10.1 | 128.2 ± 29.9 | 0.35 |
| Early antral follicle | 65.0 ± 18.0 | 50.4 ± 17.2 | 0.23 |
| Antral follicle | 8.7 ± 2.9 | 8.5 ± 3.2 | 0.92 |
| Preovulatory follicle | 6.4 ± 0.4 | 14.2 ± 11.6 | 0.17 |
| Atretic follicle | 53.1 ± 8.0 | 35.4 ± 18.1 | 0.08 |
| Total | 527.6 ± 22.3 | 913.1 ± 269.2 | <0.05 |

The data are presented as the mean ± standard deviation (SD).
\**P*<0.05 was considered to indicate statistical significance.

### The influence of ABVD withdrawal on fertility in female mice

To evaluate the effect of ABVD withdrawal on fertility in female mice, IVF experiments were performed. The results showed that the mean number of retrieved oocytes per female mouse was significantly decreased in the ABVD withdrawal group (11.0 ± 2.5 vs. 7.2 ± 1.8, *P* < 0.05). Nevertheless, there was no significant difference in the rate of cleavage or blastocyst formation (Table 3).

**Table 3.** The total number of oocytes per mouse after superovulation and blastocyst.

| Groups | Number of MII<br>oocytes per mouse | Cleavage rate (%) | No. of blastocyst (%) |
| --- | --- | --- | --- |
| Control | 11.0 ± 2.5 | 43/61 (70.5%) | 35/43 (81.4%) |
| ABVD withdrawal | 7.2 ± 1.8 | 30/41 (73.2%) | 26/30 (86.7%) |
| *P* value | < 0.05 | 0.83 | 0.75 |

Natural mating trials were then performed 4 weeks after ABVD withdrawal. Results showed that mice were fertile after ABVD treatment withdrawal. The number of offspring in the ABVD group was smaller than that in the control group (12.2 ± 1.8 vs. 9.4 ± 0.6, *P* < 0.05), but there was no significant difference in offspring weight between the ABVD group and the control group (1.6 ± 0.1 vs. 1.6 ± 0.2, *P* = 0.95) (Fig. 3).

**Fig. 3.**
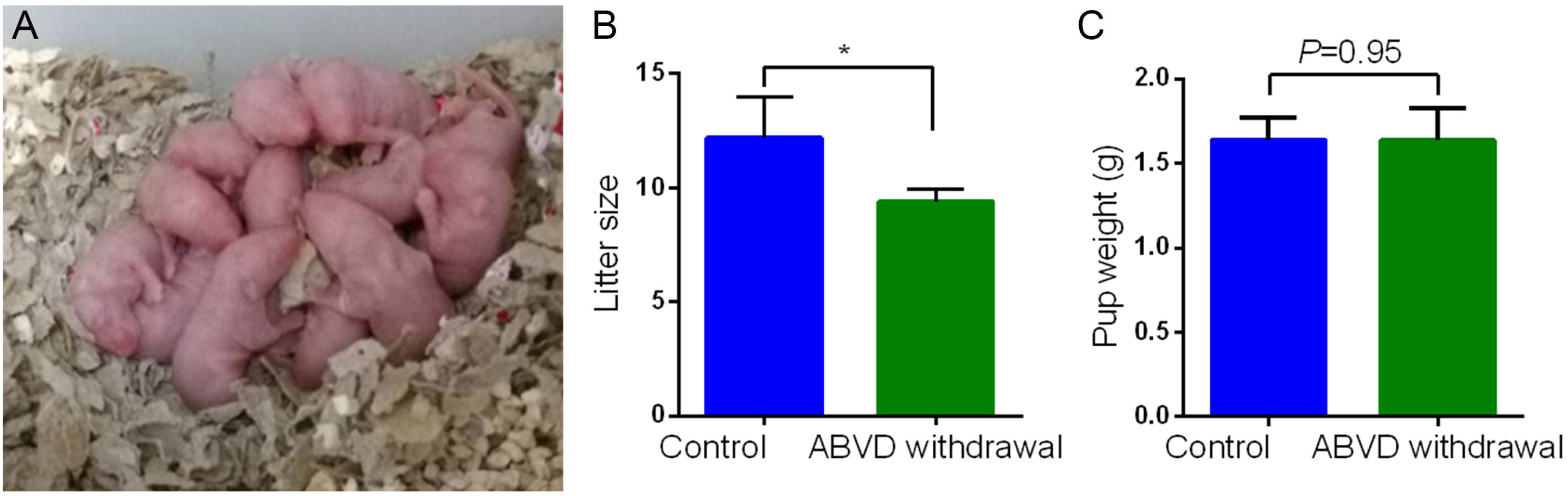
Reproductive ability was restored after ABVD treatment withdrawal. A. Mice were fertile after ABVD treatment withdrawal, and no obvious defects were found in the offspring. B. The number of pups per female decreased significantly in the ABVD withdrawal group. C. There was no difference in average pup weight between the ABVD withdrawal and control groups. The data are presented as the means ± SDs. \**P* < 0.05 compared to control.

### Transcriptome analysis of the ovaries after ABVD treatment and withdrawal in female mice

To determine the effect of ABVD treatment and withdrawal on ovarian gene expression in female mice, RNA-seq was performed. Compared with the control, a total of 751 DEGs were identified in the ABVD treatment ovaries; however, only 59 DEGs were identified in the ABVD withdrawal ovaries [-Log_2_(fold change) > 1 and *P*adj < 0.05]. Interestingly, Hspa1a and Hspa1b were upregulated in the ABVD withdrawal groups (Fig. 4A and 4B).

**Fig. 4.**
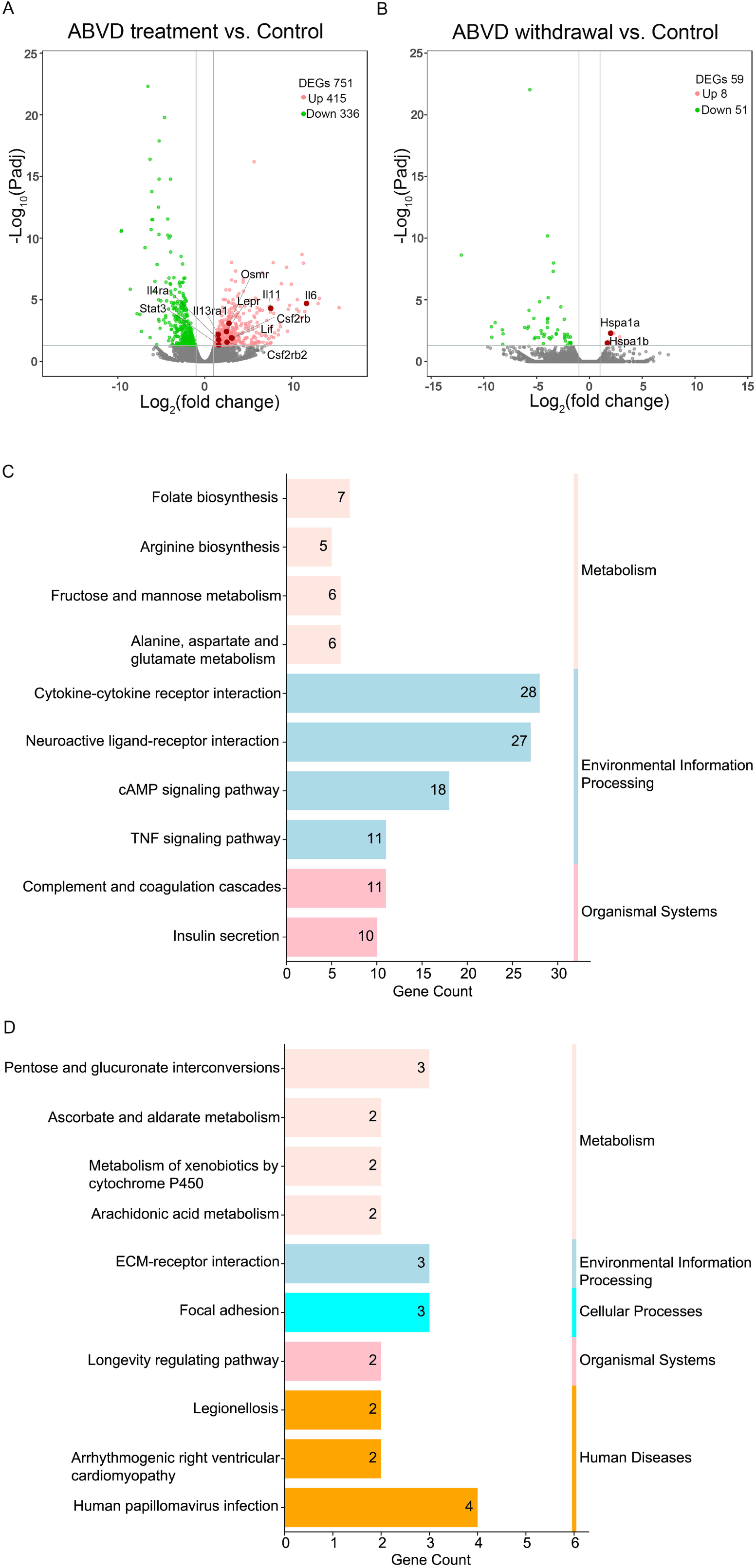
Transcriptome analysis of the effect of ABVD treatment and withdrawal on the female mouse ovaries. A. Volcano plot visualizing the DEGs in the ovary for the ABVD treatment and control groups. B. Volcano plot visualizing the DEGs in the ovary between the ABVD withdrawal and control groups. C. KEGG analysis of the DEGs in the ovaries of the ABVD treatment/control group and mapping to six functional categories. D. KEGG analysis of DEGs in the ovaries of the ABVD withdrawal/control group and mapping to five functional categories.

Then, KEGG pathway analysis of DEGs was performed. Enriched KEGG pathways were also clustered into six KEGG subsystems: metabolism, genetic information processing, environmental information processing, cellular processes, organismal systems, and human diseases. The 10 most significantly enriched pathways are shown in Fig. 4C, 4D. The results showed that most enriched pathways in the ABVD treatment group fell into the categories “environmental information processing” and “metabolism”. This means that ABVD treatment mainly affected the expression levels of genes associated with environmental information processing and metabolism in the ovary.

To further explore the effect of ABVD treatment on mouse ovaries, upregulated and downregulated DEGs were analysed separately. The results showed that pathways associated with follicular development, such as cAMP signalling, PPAR signalling, and cGMP-PKG signalling, were significantly downregulated after ABVD treatment; however, the JAK/STAT pathway, which is involved in the inhibition of primordial follicle activation and apoptosis, was upregulated (Fig. 5A and 5B). Heat map analysis demonstrated that at least 8 genes involved in cAMP signaling pathway were down-regulated in ovaries of ABVD treatment mice. And at least 12 genes involved in MAPK signaling pathway and Jak-STAT signaling pathway were up-regulated in ovaries of ABVD treatment mice, respectively (Fig. S2).

**Fig. 5.**
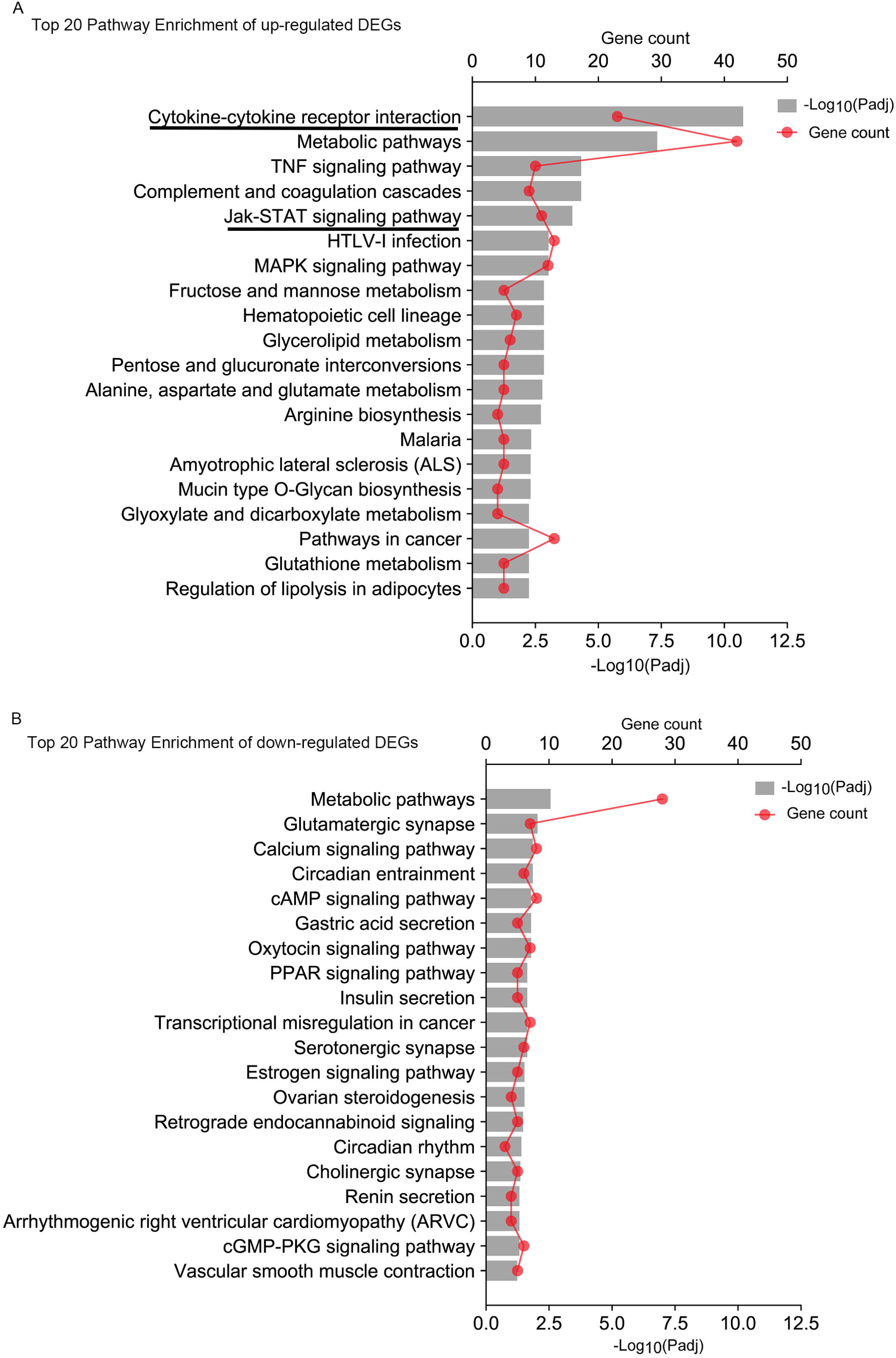
KEGG pathway enrichment of upregulated and downregulated DEGs in ABVD-treated ovaries. A. Top 20 pathways enriched in the upregulated DEGs in ABVD-treated ovaries. B. Top 20 pathways enriched in the downregulated DEGs in ABVD-treated ovaries. The top axis is the enrichment gene count, and the bottom axis is -log_10_ (adjusted p values).

Ligand-receptor binding leads to activation of the JAK/STAT pathway. In detail, we found that the ligand (Il11, Lif, and Il6), receptor (Lepr, Osmr, Csf2rb, Il13ra1, Il4ra, and Csf2rb2), and signal transducer and activator of transcription (Stat3) genes of the JAK/STAT pathway were significantly upregulated after ABVD treatment (Fig. 4A, Fig. S3 and Fig. S4).

## Discussion

The majority of women experience amenorrhea after ABVD treatment, and serum AMH levels fall significantly[18]. However, after the end of chemotherapy, most women return to regular menstrual cycles, and AMH concentrations are restored to pre-treatment values[17]. Our results showed that the mice lost their regular estrous cycles soon after ABVD treatment and that body weight increased much more slowly. The gross ovarian weight, serum AMH and number of ovarian primordial follicles decreased dramatically compared with those of the control after 4 cycles of ABVD treatment. This mouse model simulated a typical situation: ABVD treatment in young women with Hodgkin lymphoma.

Previous studies have showed that ovarian cortical tissue from women treated with ABVD contains a higher density of nongrowing follicles (NGFs). They speculated that the total number of NGFs was also increased in women treated with ABVD[19]. Our research provides direct evidence that compared to those of the control group, serum AMH levels and the total number of NGFs were increased after ABVD withdrawal. We also found that the mean numbers of retrieved oocytes and offspring were significantly decreased in the ABVD withdrawal group. However, the rate of cleavage and blastocyst formation and offspring weight were similarly unaffected. Previous studies found similar results. Hodgson et al. found no evidence of significant impairment of fertility in females who had received ABVD compared to controls[15]. However, tissue fragments from ABVD-treated ovaries resulted in a reduction in the in vitro developmental potential, analogous to that observed in ovaries from prepubertal girls[31].

The pool of NGFs is used to describe the remaining ovarian reserve[32]. Notably, the results of the present study demonstrated that the number and proportion of NGFs in mouse ovaries were increased four weeks after ABVD discontinuation, suggesting that the ovarian reserve was elevated. The molecular mechanisms of increases in NGFs after ABVD treatment are poorly documented. Previous studies suggested that the reason may be that ABVD treatment resulted in the formation of new follicles[19]. However, evidence for this has not been provided to date[9, 33]. In this study, transcriptome sequencing revealed that pathways associated with follicular development, such as cAMP signalling, PPAR signalling, and cGMP-PKG signalling, were significantly downregulated after ABVD treatment[34, 35]. Moreover, the JAK/STAT pathway, which is involved in the inhibition of primordial follicle activation and apoptosis, was upregulated[33].

The JAK/STAT pathway is a classical signal transduction pathway whose activation relies on ligand binding to the receptor. We found that the ligand (Il11, Lif, and Il6), receptor (Lepr, Osmr, Csf2rb, Il13ra1, Il4ra, and Csf2rb2) and signal transducer and activator of transcription (Stat3) genes of the JAK/STAT pathway were significantly upregulated after ABVD treatment. A possible reason is that these ligands and receptors are stimulated after ABVD treatment, and pathways involved in the inhibition of primordial follicle activation and apoptosis are activated. This study investigated the transcriptomic profiles of whole ovary to reveal the effect of ABVD treatment on ovarian gene expression, which could not reflect what fraction of ovary were affected after ABVD treatment. Spatial transcriptomics should be performed to get more accurate results in the future.

Adriamycin is a first-line anticancer agent and is widely applied in the clinic. A previous study reported that adriamycin compromised fertility by damaging ovarian follicles and showed dose-dependent toxicity in mice. Adriamycin displayed notable ovarian toxicity when administered at 0.4 mg/kg[36]. However, in our research, mice were treated with adriamycin 8.3 mg/kg administered weekly for 4 cycles as a polychemotherapy regimen and did not exhibit evident ovarian toxicity. Some studies have also reported that dacarbazine aggravated the reduction in ovarian reserve through depletion of primordial follicles in mice[37]. Surprisingly, it seems that combined ABVD treatment has a beneficial effect on the ovarian reserve. This phenomenon may be ascribed to the synergistic effects of ABVD.

Despite the lower gonadotoxic risk of this treatment, the necessity of fertility preservation before ABVD treatment remains controversial. This study provides important evidence on the reproductive toxicity of ABVD chemotherapy. The ovarian follicular reserve and reproductive capacity were able to recover after ABVD withdrawal. Fertility preservation for young female patients who receive ABVD therapy alone is not desired. Additionally, ABVD chemotherapy causes significant reductions in fertility, and some patients still need more aggressive therapy[8]. All these issues also need to be considered when performing fertility preservation counselling. The mechanism underlying the enhancement of ovarian reserve after ABVD chemotherapy needs further clarification. Women with Hodgkin’s lymphoma are generally performed ABVD therapy in clinical practice. Due to the mice model used here lacking a corresponding disease, the outcomes may be affected to some extent. Mice model with Hodgkin’s lymphoma is needed for more research in the future.

## Conclusions

In summary, ABVD chemotherapy may have an impact on ovarian function. However, the ovarian function and fertility can be partially recovered after ABVD discontinuation, and the ovarian reserve was increased instead compared with the control. ABVD chemotherapy for a short time could suppresses the activation and apoptosis of primordial follicles through the JAK/STAT pathway and then improve the ovarian reserve.

## List of abbreviations

ABVD: adriamycin, bleomycin, vinblastine and dacarbazine
HL: hodgkin lymphoma
DOR: diminished ovarian reserve
POF: premature ovarian failure
AMH: anti-Mullerian hormone
H&E: haematoxylin and eosin
IVF: in vitro fertilization
PMSG: pregnant mare serum gonadotropin
hCG: human chorionic gonadotrophin
COCs: cumulus oocyte complexes
HTF: human tubal fluid
KEGG: Kyoto Encyclopedia of Genes and Genomes
NGFs: nongrowing follicles.

## Declarations

### Ethics approval and consent to participate

This study was approved by the Ethics Committee of Shanghai Tenth People’s Hospital and conducted in accordance with Tongji University animal research requirements.

### Consent for publication

Not applicable.

### Availability of data and materials

Original data and materials are available from the corresponding author on reasonable request.

### Authors’ contributions

Y.B.L. contributed to the study design, writing and critical discussion. Y.B.L and X.M.L. performed experiments execution and data analysis. R.C.C. and B.T. contributed to the experiments execution and manuscript comment. X.M.L. and Y.B.L. contributed to the drafting the manuscript. All authors read and approved the final manuscript.

## Acknowledgements

Not applicable.

## FIGURE CAPTIONS

**Supplementary Fig. 1** Comparison of the number of primordial follicles after ABVD treatment and withdrawal.

**Supplementary Fig. 2** Heat map analysis of the differentially expressed genes involved in cAMP signalling pathway, MAPK signalling and Jak-STAT signalling pathway, respectively. A. Heat map analysis of the differentially expressed genes involved in cAMP signalling pathway. B. Heat map analysis of the differentially expressed genes involved in MAPK signalling pathway. C. Heat map analysis of the differentially expressed genes involved in Jak-STAT signalling pathway.

**Supplementary Fig. 3** The JAK-STAT signalling pathway, showing upregulated genes in the ovaries of ABVD-treated mice versus control mice. Red marks indicate the genes with significantly upregulated expression.

**Supplementary Fig. 4** The cytokine–cytokine receptor interaction pathway showing DEGs in the ovaries of ABVD-treated mice versus control mice. Red boxes: upregulated genes; green boxes: downregulated genes.

