## Supplementary material for "Fertility recovery after combined adriamycin, bleomycin, vinblastine and dacarbazine chemotherapy in mice": S1: Supplementary_Material.pdf

**Running Title:** Fertility recovery following combined ABVD chemotherapy

### *Supplementary Material*

Supplementary Figures

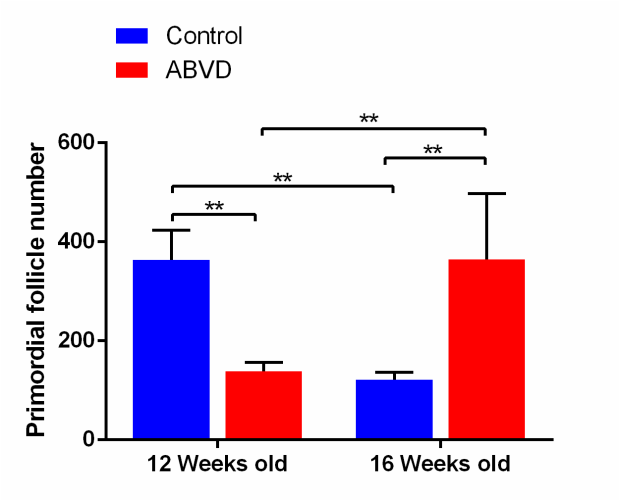


**Supplementary Fig. 1** Comparison of the number of primordial follicles at after ABVD treatment and withdrawal.

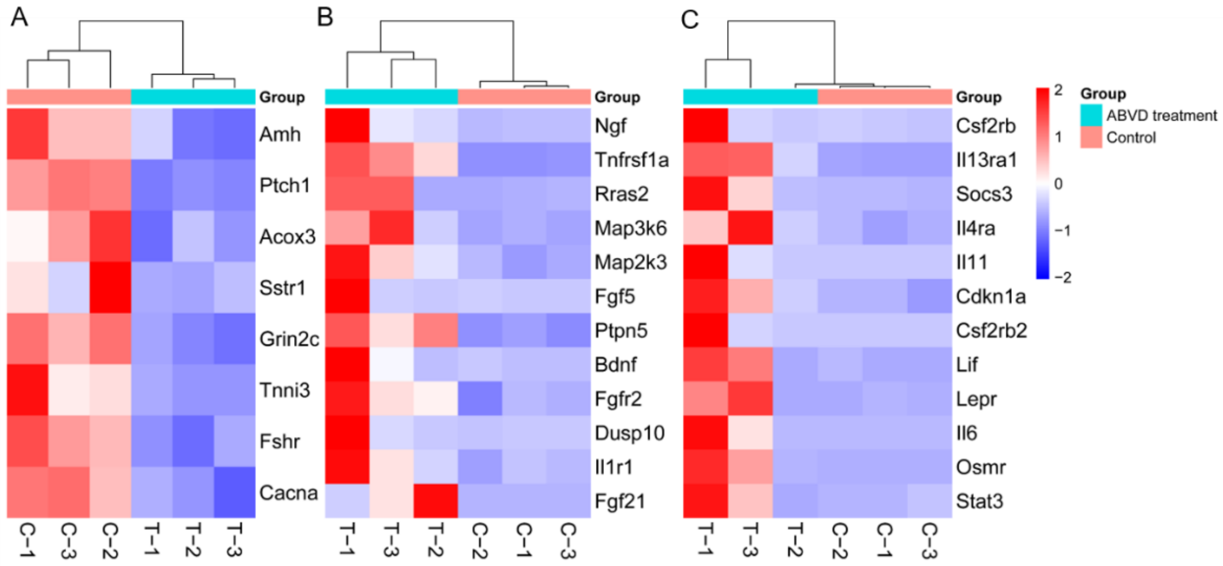


**Supplementary Fig. 2** Heat map analysis of the differentially expressed genes involved in cAMP signaling pathway, MAPK signaling and Jak-STAT signaling pathway, respectively. A. Heat map analysis of the differentially expressed genes involved in cAMP signaling pathway. B. Heat map analysis of the differentially expressed genes involved in MAPK signaling pathway. C. Heat map analysis of the differentially expressed genes involved in Jak-STAT signaling pathway.

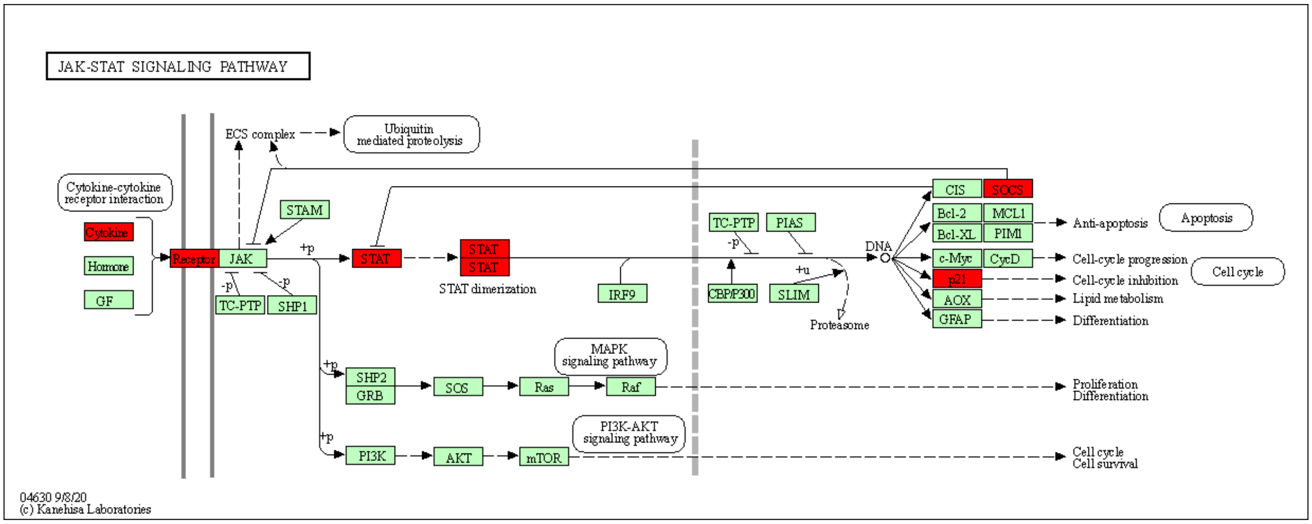


**Supplementary Fig. 3** The JAK-STAT signaling pathway, showing upregulated genes in the ovaries of ABVD-treated mice versus control mice. Red marks indicate the genes with significantly upregulated expression.

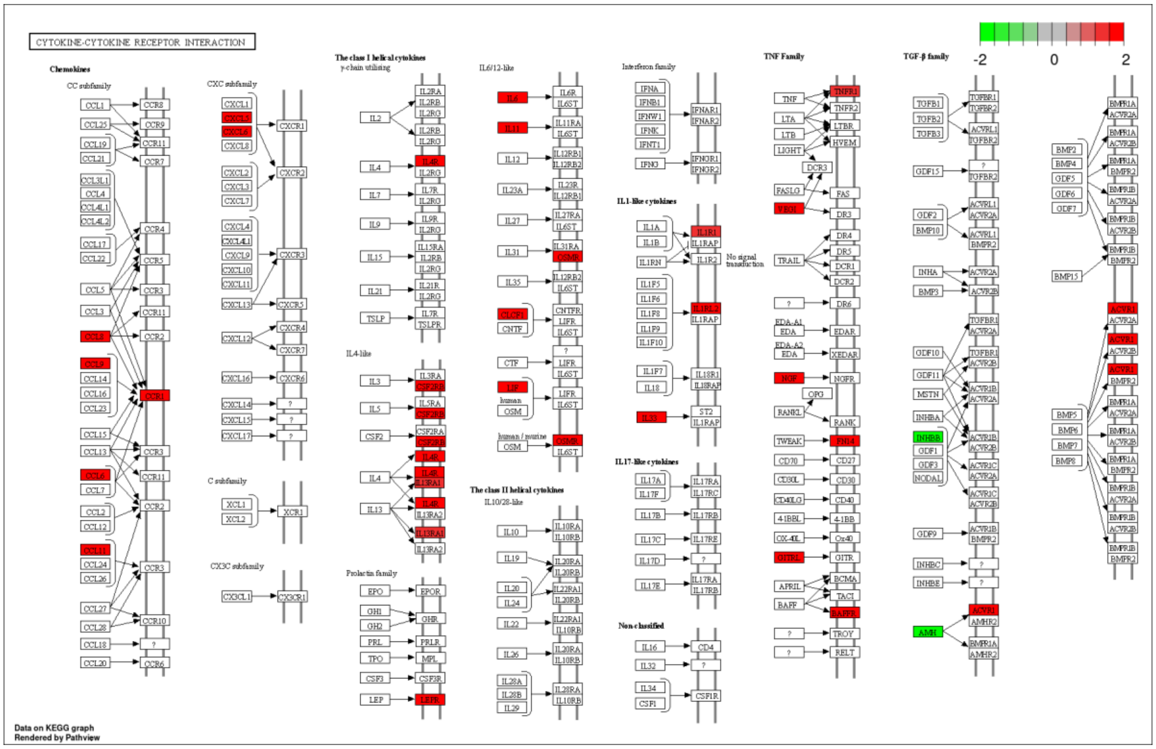


**Supplementary Fig. 4.** The cytokine–cytokine receptor interaction pathway showing DEGs in the ovaries of ABVD-treated mice versus control mice. Red boxes: upregulated genes; green boxes: downregulated genes.
